# Tuning a mechanical model to biological reality: A case study in the LaMSA system of the trap-jaw ant *Strumigenys*

**DOI:** 10.1101/2024.02.15.580213

**Authors:** Philip S L Anderson, Justin F Jorge, Stephanie B Crofts, Jackson T Castro, Rosalie L Didcock, Andrés Cook, Fredrick J Larabee, Mark Ilton

## Abstract

Understanding the relationship between morphology and movement in biomechanical systems, particularly those composed of multiple complex elements, presents challenges due to the nonlinear nature of the interaction between components. This study focuses on the mandibular closing mechanisms in ants, specifically comparing muscle-driven actuation (MDA) and latch-mediated spring actuation (LaMSA) in the genus *Strumigenys*. Analyzing 3D structural data from diverse *Strumigenys* species, we employ mathematical models for both LaMSA and MDA systems. Our findings reveal distinct patterns of mechanical sensitivity between the two models, with sensitivity varying across kinematic output metrics. We explore the performance transition between MDA and LaMSA systems by incorporating biological data and correlations between morphological parameters into the models. In these models tuned specifically to *Strumigenys*, we find the LaMSA mechanism outperforms MDA at small relative mandible mass. Notably, the location and abruptness of the performance transition differs among various kinematic performance metrics. Overall, this work contributes a novel approach to understanding form-function relationships in complex biomechanical systems. By using morphological data to calibrate a general biomechanical model for a particular group, it strikes a balance between simplicity and specificity and allows for conclusions that are uniquely tuned to the morphological characteristics of the group.

## INTRODUCTION

The relationship between biological morphology and the corresponding motion or kinematics produced by said morphology is often nonlinear, especially in biomechanical systems comprised of multiple elements in complex configurations (Koehl 1996; Wainwright et al. 2005). These complicated configurations emerge because of the rich evolutionary histories of these biomechanical systems and are a natural consequence of organisms using their morphology for numerous functions (locomotion, feeding, defense, etc.). The non-linear nature of these biomechanical systems also leads to a conundrum for researchers interested in teasing apart the relationships between morphology and function: Seemingly small changes in biological morphology can cause large, unintuitive changes in biomechanical performance (Koehl 1996; Wainwright et al. 2005). Identifying these relationships between morphological change and biomechanical consequence is made difficult by the very nature of these complex systems, especially when trying to compare systems across taxa. Here, we utilize a set of novel biomechanical modeling methods designed to address the inherent non-linearity of biomechanical systems to understand the form-function relationship across two distinct mandibular closing mechanisms in ants.

In a seminal work, Koehl (1996) defined specific aspects of multi-part biological systems that complicate the relationship between morphology and mechanical performance. First, across the range of variation in a morphological parameter, a particular change can have a minimal effect or can have large effect on the mechanical output (performance) (Koehl 1996). This phenomenon, termed mechanical sensitivity, has been demonstrated in fish oral jaws, mantis shrimp appendages, and lizard jaws (P. S.L. Anderson and Patek 2015; Hu, Nelson-Maney, and Anderson 2017; Baumgart and Anderson 2018; Cruz et al. 2021). Second, the various morphological parameters may not be independent of each other but may vary in a systematic way such that the mechanical output of a system is not simply the additive effect of its morphological inputs (Koehl 1996). The interdependence of biological morphologies has been well documented and hypothesized to be caused by structural, developmental, or even genetic integration (Breuker, Debat, and Klingenberg 2006). Third, morphologically distinct configurations of a biomechanical system can be ‘functionally equivalent’, resulting in equivalent performance. This phenomenon, also called many-to-one mapping (Wainwright et al. 2005), has been widely studied in biomechanical systems (Alfaro, Bolnick, and Wainwright 2004; 2005). Finally, biomechanical systems in nature, may have multiple biologically-relevant kinematic output metrics. In this case, each of these metrics may respond differently to the same underlying variation in morphological inputs (Baumgart and Anderson 2018).

The complex relationship between morphology and biomechanical performance is particularly important when trying to understand the evolution of these systems. For example, mantis shrimp, trap-jaw ants, and jumping insects have all shown evolutionary transitions from a direct muscle-driven actuation (MDA) of movement to using latch-mediated spring actuation (LaMSA) (Sutton et al. 2022). Biomechanical modeling has been used to understand how both morphology and performance can change across these transitions, and how those changes relate to each other. A primary determinant of the benefit of a LaMSA mechanism is the size of the animal. For a given system, a LaMSA mechanism achieves higher performance than the MDA mechanism only below a certain mass threshold (Galantis and Woledge 2003; Ilton et al. 2018; Cook et al. 2022). To date, modeling of this performance transition has included general assumptions about size-scaling of the morphological parameters that may not be relevant to specific biological systems. In this work, we explore we explore both the morphological and performance transitions between MDA and LaMSA systems within the ant Genus *Strumigenys*.

LaMSA-driven trap-jaw mechanisms have evolved multiple times across ants, with each evolutionary origin converging on a general schema (Fig. 1): a muscle loads potential energy into a spring element (i.e., apodeme and/or cuticle) attached to the mandibles, while a latch holds the mandible in place (Larabee and Suarez 2014). The nature of this latch varies, but in all cases the jaws snap shut at extreme velocities when that energy is released (Gibson 2021), reaching up to 60 m/s in some species (Patek et al. 2006; Sutton et al. 2022). LaMSA allows trap-jaw ants to actively hunt evasive prey such as springtails, leaf-litter arthropods with fast escape behaviors, whereas species with MDA mandibles utilize hunting strategies based on stealth or chemical deception (Brown Jr and Wilson 1959; Dejean 1985; Masuko 1984). *Strumigenys* is a globally distributed, hyperdiverse genus of ants (800+ species) with a wide diversity of mandible types including up to 10 independent origins of the LaMSA trap-jaw system (Booher et al. 2021). Mirroring their morphological diversity, *Strumigenys* species also display diverse mandible performance, including variation in velocity, acceleration, and power density (Booher et al. 2021; Gibson 2021). The wide diversity of both LaMSA and MDA morphology and mechanisms makes *Strumigenys* an excellent clade for exploring variation in biomechanical models.

**Figure 1:**
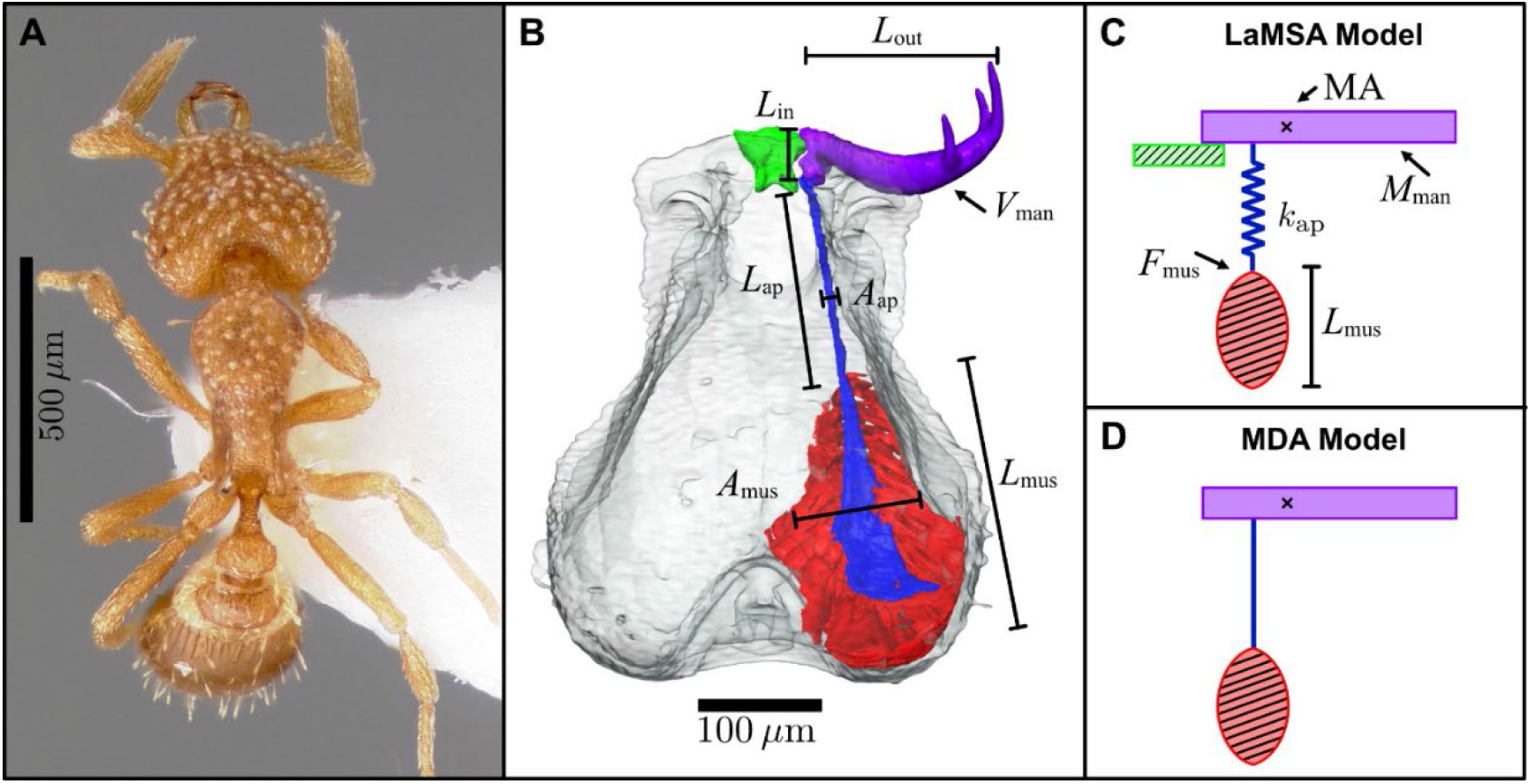
Morphological features, parameter definitions, and model schematics used in this study. **A**. Dorsal view of *Strumigenys emmae* (photographer: April Nobile, specimen: CASENT0005894 from www.antweb.org). **B**. MicroCT rendering of *Strumigenys emmae* with different colors representing key morphological features for mandible closure: red - closer muscle; blue - apodeme; purple - mandible; and green - labrum. We measure mandible volume (*V*_man_), and the in-lever (*L*_in_) and out-lever (*L*_out_) arms of the closing mandible. For the apodeme, we measure its length (*L*_ap_) and its cross-sectional area (*A*_ap_). We measure the length of the closer muscle (*L*_mus_), the area of muscle attachment to the head capsule (*A*_mus_), and the median angle of muscle attachment for 10 muscle fiber bundles. **C**. These morphological measurements are used to compute five focal parameters used in the Latch-Mediated Spring Acutation (LaMSA) model: the mechanical advantage (MA) around a fixed rotation axis “x”, the mass of the mandible (*M*_man_), the spring constant of the apodeme (*k*_ap_), the maximum muscle force (*F*_mus_), and muscle length. Using these parameters, the LaMSA model computes the mandible dynamics assuming that the closer muscle contracts isometrically to load the apodeme, while a latch (shown in green) holds the mandible in place. After loading is complete, the latch is removed from the system and the apodeme pulls on the mandible to drive a spring-driven movement. **D**. These same five parameters are used in the direct Muscle-Driven Actuation (MDA) model except the apodeme is assumed to be rigid (*k*_ap_ → ∞) and there is no latch. The closer muscle contraction directly

To explore the mechanical sensitivity within both MDA and LaMSA mechanisms in ant mandibles and the performance transition between them, we incorporate 3D structural data from select *Strumigenys* species (both trap-jaw and non-trap-jaw forms) into two mathematical models for mandible actuation: one based on a trap-jaw LaMSA system and the other on an MDA system driven by muscles (Fig. 1). Using these models, we address the following questions: 1) Mechanical Sensitivity: Are patterns of mechanical sensitivity similar between LaMSA and MDA systems with the same underlying morphology across different kinematic output metrics? 2) Performance Transition: When we include interdependence between morphological inputs in our LaMSA and MDA models, does the transition between MDA and LaMSA systems coincide with observed relative size differences between trap-jaw and non-trap-jaw *Strumigenys* species? Is the performance transition qualitatively similar across different metrics of fast kinematic performance? By answering these questions, we identify areas in the *Strumigenys* morphospace and scaling relationships that can be further probed in future experiments.

## METHODS

### Animal collection and husbandry

Seven of the 12 *Strumigenys* species were obtained from colonies collected by the Suarez lab (School of Integrative Biology, University of Illinois Urbana-Champaign). *Strumigenys rostrata* colonies were collected from rotting logs at Brownfield woods, Illinois (40.14461°N, 80.1657°W ±100m) in May of 2016; *Strumigenys denticulata* colonies were collected from rotting logs at Nouragues Ecological Field station, French Guiana (3.982411°N, 52.563872°W ±200m) in March of 2016; a *Strumigenys louisianae* colony was collected from a rotting log at Magnolia Springs State park, Georgia (32.88068°N, 81.95612°W ±500m) in August of 2015; *Strumigenys eggersi* colonies were collected from fallen acorns at Archbold Biological Station, Florida (27.18285°N, 81.35208°W ±1km) in August of 2015 and May of 2016; colonies of *Strumigenys auctidens* and *Strumigenys decipula* were collected from rotting twigs in leaf litter at the Amazon Conservatory for Tropical Studies field station (ACTS) in Loreto, Peru in July of 2017 and 2018 (3°14’60.00”S, 72°54’36.00”W ±1km); several workers of *Strumigenys trinidadensis* were collected at ACTS in July of 2017 by sifting leaf litter through a winkler extractor with a cup containing moist paper towels placed at the bottom rather than preservative. All ants were collected legally with the appropriate permits.

All colonies or collections of individual workers were kept in 10.5 cm x 10.5 cm x 3 cm fluon coated plastic sandwich containers with pieces of their source substrate when possible, or a layer of moist vermiculite otherwise, and stored in a USDA certified quarantine facility continuously kept at approximately 25°C, 50% relative humidity and on a 12h day-night cycle between filming sessions. All were fed ad libitum laboratory raised temperate springtails purchased from Genesis Exotics (Houston, TX) or Josh’s Frogs (Owosso, MI) while housed in the lab.

### Specimen preparation and microCT

For each species, one specimen was fixed for 24h in 70% alcoholic Bouin’s solution to preserve internal structures, after which they were washed twice in 70% ethanol and dehydrated to 100% ethanol. Prior to scanning, each specimen was stained overnight in a 1% iodine in 100% ethanol before being washed twice in 100% ethanol and dried using a 931.GL Supercritical AutoSamdri Critical Point Dryer (Tousimis Research Corporation, Rockville, MD). This approach improved the contrast between the ants’ tissues and background. Once dried, whole specimens were either placed into a pipette tip or the specimen’s head was carefully glued to the tip of a toothpick at the vertex of the head using superglue. Specimens were scanned with a Xradia MicroXCT-400 scanner (Carl Zeiss, Oberkochen, Germany) using voxel sizes 0.912–1.51 µm^3^. Additionally, we used previously published scans of *Strumigenys depressiceps, Strumigenys elongata, Strumigenys emmae, Strumigenys faurei and Strumigenys simoni* (Booher et al. 2021).

Anatomical components in the microCT scans were measured using Avizo lite and the digital calipers in Geomagic (3D Systems). We isolated and segmented the structures involved in the mandible movement for each species, isolating the right mandible and associated apodeme for each specimen scanned. As detailed in the next section, we also isolated muscle components to estimate the force input to the mandible.

### Morphological measurements

Using measurements of the microCT scans (Fig. 1B), we calculated five components used in the LaMSA and MDA models (Fig. 1C-D). *Mechanical advantage* (MA) was calculated based on linear measurements of the mandibular lever arms involved in jaw closing (Fig. 1B). *Mandible mass* was calculated from the mandible volume using a density of 1.3 kg/m^3^ previously measured for similar tissue (Vincent and Wegst 2004). *Apodeme spring constant* (*k*_ap_) was calculated using the length (*L*_ap_) and maximum cross-sectional area (*A*_ap_) of the closer muscle apodeme at the distal-most point of muscle attachment (Fig. 1B), where 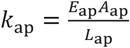. The apodeme modulus (*E*_ap_) was set as 11 GPa based on previous measurements of similar cuticle (Vincent and Wegst 2004). *Closer muscle length* (*L*_mus_) and *maximum closer muscle force* (*F*_mus_) were calculated using a previously-established equation (Alexander 1969) as follows

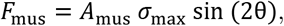

where *A*_mus_ is the measured area of muscle attachment to the head capsule. The median muscle fiber angle (θ) was calculated from measurements of isolated individual muscle fiber bundles distributed throughout the volume of the closer muscle. Maximum muscle fiber stress (*σ*_max_), 300 kN/m^2^, was based on previous research (Gronenberg 1996).

Non-dimensionalization of model parameters models, which makes computation more efficient and data visualization easier. Non dimensionalization has been performed in prior modeling work on both LaMSA and MDA systems (Galantis and Woledge 2003; Ilton et al. 2018; Labonte 2023). For nondimensionalizing the dynamics of a mechanical system, we needed to choose three characteristic quantities: a characteristic mass *M*_*c*_, a characteristic length *L*_*c*_, and a characteristic time *T*_*c*_ (for a survey of the principles of nondimensionalization, see Langtangen and Pedersen 2016). Here we constructed a characteristic length, mass, and time based on the properties of the apodeme because the role of the apodeme spring is central in this work. To construct these three characteristic quantities, three independent properties of the apodeme were used: the apodeme length *L*_*c*_, the apodeme density *ρ*_ap_, and the apodeme modulus *E*_ap_. Combinations of *L*_*c*_, *ρ*_ap_, and *E*_ap_ were then used to create the characteristic scaling quantities *M*_*c*_, *L*_*c*_, and *T*_*c*_ as follows. The length of the apodeme was used directly to set a characteristic size scale *L*_c_ = *L*_ap_, consistent with that chosen in previous work (Ilton et al. 2019). To construct a characteristic mass, we used the characteristic length and the density of the apodeme *ρ*_ap_ because we assume that there are relatively small variations in apodeme density across the different species. We used *ρ*_ap=_1.3 kg/m^3^ to match the previously measured values for sclerotised cuticle (Vincent and Wegst 2004) (small changes to our assumed value of *ρ*_ap_ within a reasonable range of 1.0-1.3 kg/m^3^ does not affect the results of this work – for example, a 10% difference in the assumed *ρ*_ap_ results in shifts of the biological data in Figures 3-4 that are smaller than the individual data points). This density was used with the characteristic length to construct a characteristic mass-scale as *M*_c_ = *ρ*_ap_ *L*_c_^3^. Finally, a characteristic timescale is set by the propagation of elastic waves through the apodeme based on its modulus, *E*_ap_, as *T*_c_ = *L*_c_ (*ρ*_ap_/*E*_ap_)^1/2^ (Ilton et al. 2019).

Based on these characteristic quantities, we normalized all the calculations and measurements. For example, the maximum isometric muscle force, which was calculated based on the estimated muscle physiological cross-sectional area, was normalized (non-dimensionalized) by a characteristic force 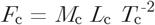, which was constructed from *M*_c_, *L*_c_, and *T*_c_ to give units of force. Except where otherwise noted, all quantities in this manuscript are reported in terms of normalized units. Consequently, from our simulations in the next section, maximum mandible velocity during a strike, or “take-off” velocity *v*_to_, is reported in dimensionless units as a multiple of the characteristic velocity *V*_c_ = *L*_c_ / *T*_c_. Tables 1 and 2 summarize the measured morphological characteristics for each species and their normalized values that serve as inputs to the models.

**Table 1:** Dataset of the morphological parameters measure from microCT scans of *Strumigenys* ants.

|  | Mandible |  |  | Closer Muscle |  |  | Apodeme |  |
| --- | --- | --- | --- | --- | --- | --- | --- | --- |
| | $L_{in}$<br>($\mu m$) | $L_{out}$<br>($\mu m$) | $V_{man}$<br>($\mu m^3$) | $L_{mus}$<br>($\mu m$) | $A_{mus}$<br>($\mu m^2$) | $\theta$<br>($^\circ$) | $L_{ap}$<br>($\mu m$) | $A_{ap}$<br>($\mu m^2$) |
| **Trap-jaw species** |  |  |  |  |  |  |  |  |
| *S. emmae* | 26.1 | 140 | 73553 | 243 | 61933 | 44.3 | 352 | 83.6 |
| *S. faurei* | 24.5 | 253 | 269232 | 357 | 92022 | 39.5 | 389 | 63.5 |
| *S. elongata* | 41.6 | 382 | 516780 | 423 | 158009 | 38.2 | 480 | 97.8 |
| *S. trinidadensis* | 58.5 | 550 | 1341401 | 522 | 236344 | 41.3 | 586 | 212.9 |
| *S. auditens* | 58.7 | 255 | 227127 | 366 | 73276 | 27.2 | 321 | 73.6 |
| *S. eggersi* | 31.9 | 276 | 133730 | 270 | 55430 | 35.2 | 253 | 141.5 |
| *S. louisianae* | 30.9 | 289 | 317497 | 363 | 108344 | 36.5 | 327 | 73.3 |
| *S. decipula* | 65.8 | 318 | 512076 | 395 | 132117 | 39.2 | 365 | 182.0 |
| *S. denticulata* | 64.8 | 369 | 356419 | 283 | 57455 | 43.9 | 226 | 56.8 |
| **Non-trap jaw species** |  |  |  |  |  |  |  |  |
| *S. rostratus* | 47.6 | 261 | 352020 | 506 | 80938 | 16.1 | 200 | 230.7 |
| *S. depressiceps* | 32.7 | 258 | 320244 | 501 | 73553 | 14.8 | 105 | 168.1 |
| *S. simoni* | 35.1 | 272 | 396819 | 402 | 108715 | 18.2 | 65 | 291.6 |

**Table 2:**
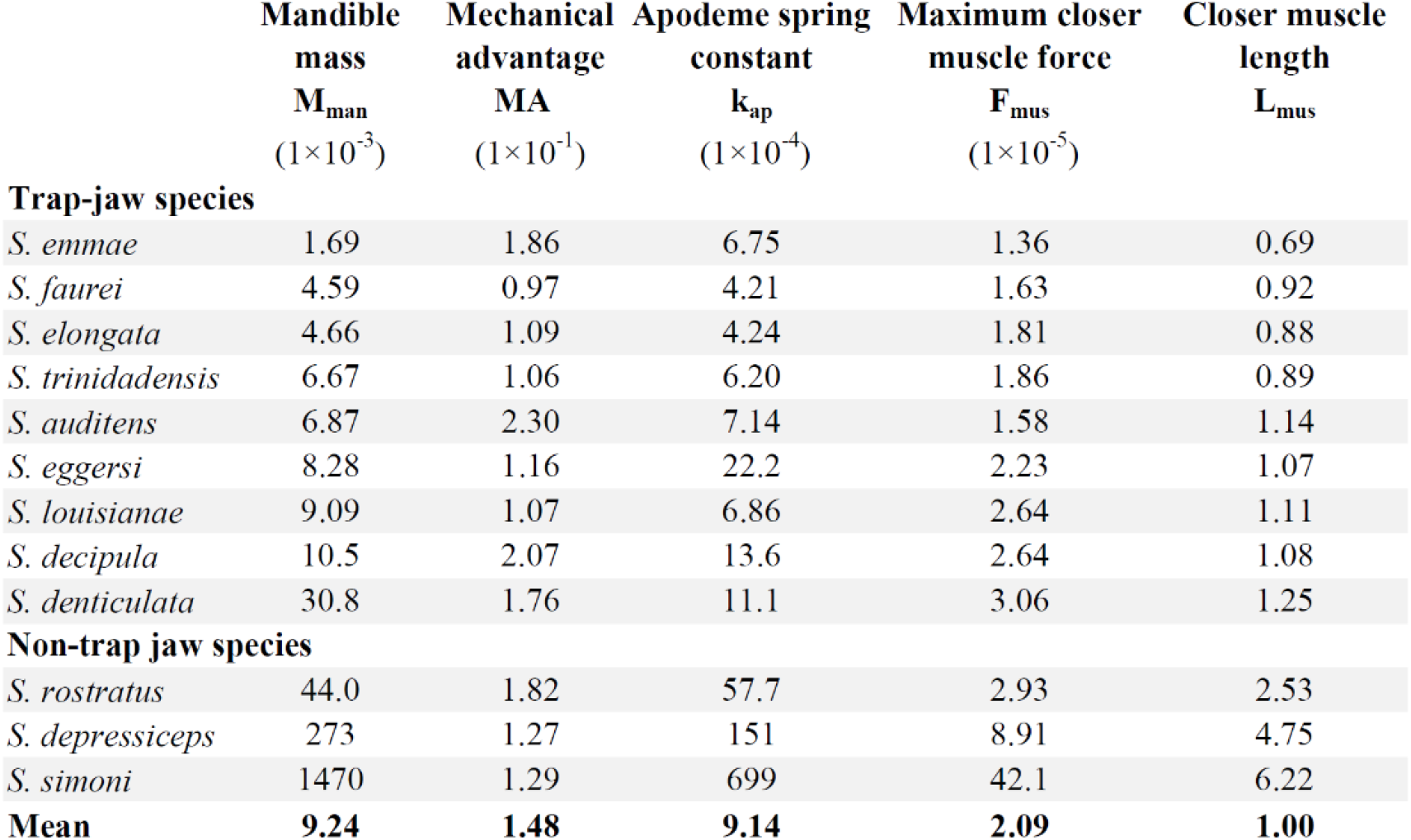
Nondimensionalized focal input parameters. Using the data from Table 1, we calculated five dimensionless focal input parameters (see Methods: Morphological measurements).

**Table 3:** Relationships between parameters used as inputs to the models. Using the data from Table 2, we created best-fit relationships between the input parameters. These relationships were used to in the models to calculate the results in Figs. 3-4.

| Model input<br>parameter | Relationship to $M_{\text{man}}$ used in the models |
| --- | --- |
| $F_{\text{mus}}$ | $F_{\text{mus}} = 0.0028 \cdot (M_{\text{man}}^{0.33}) + 2.3 \times 10^{-5}$ |
| $k_{\text{ap}}$ | $k_{\text{ap}} = 0.05 \cdot M_{\text{man}}^{0.87}$ |
| $L_{\text{mus}}$ | $L_{\text{mus}} = 5.8 \cdot M_{\text{man}}^{0.33}$ |
| MA | MA = 0.15 (constant) |

### Mathematical models

We incorporated these calculations into a previously-published dynamical simulation (Cook et al. 2022). This simulation compares the dynamics of a muscle driving a load mass in two ways: 1) the muscle directly actuates the mass (MDA model) or 2) the same muscle is used to load elastic energy into a spring-latch system and the mass is then driven by the spring (LaMSA model).

We use components in the models that reflect the *Strumigenys* morphology. For the load mass, we use a lever arm that rotates to model mandible motion (Figure 1C-D). The muscle in the model represents the large closer (adductor) muscles in the head capsule of *Strumigenys* that load elastic energy into the apodeme (spring), which connects medially to the mandible base (Booher et al. 2021) (Figure 1). The latch in the LaMSA model represents the contact between the basal mandible process and labrum in *Strumigenys* (Booher et al. 2021). For the MDA model, the large closer muscles are used to directly actuate the mandible, and the apodeme and basal mandible process are ignored.

We ran both LaMSA and MDA simulations varying the five focal model parameters over their biological range (Table 2) to calculate four kinematic output metrics of the mandibles: maximum power (*P*_max_), maximum kinetic energy (*K*_max_), take-off velocity (*v*_to_), and maximum acceleration (a_max_).

### Model sensitivity analysis

Sensitivity analyses are used in a wide range of disciplines to identify the primary factors influencing the output of a system (Frey and Patil 2002). To analyze the sensitivity of the kinematic output of each of our models to different morphological input parameters, we take a similar approach to (Carmichael and Sandu 1997) and compute the normalized sensitivity coefficients in which the relative change of output is calculated as a function of a small relative change in each input variable, holding all other variables fixed. For example, the sensitivity of the take-off velocity with respect to mandible mass is computed by normalizing the partial derivative,

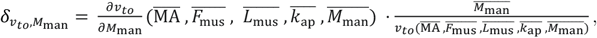

where the horizontal bars indicate an evaluation of the model at the average value of each parameter. The dimensionless sensitivity of take-off velocity for each of the five primary input variables 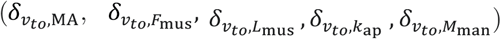 was similarly calculated for both LaMSA and MDA systems, evaluating models at the average parameter values measured from the morphological data (i.e. the normalized parameter values reported in the bottom row of Table 2).

### Comparison of LaMSA and MDA mandible movement

To compare the kinematics of a LaMSA system and an MDA system, we simulated the dynamics of both systems with the same set of muscle and load mass input parameters. We quantitatively compare LaMSA and MDA movement through calculation of a “LaMSA Ratio” for each of the kinematic output metrics. For *v*_to_, the LaMSA Ratio is defined by the ratio of take-off velocity for the LaMSA system to the MDA system, *R*_*v*to_= *v*_to_,LaMSA/*v*_to_,MDA. Similarly, we calculated *R*_*K*max_, *R*_*P*max_ and R_*a*max_. *R* = 1 corresponds to equal performance between a LaMSA system and a comparable MDA system, while *R* > 1 corresponds to a better performance of the LaMSA system.

The variation of each LaMSA ratio across a variation in the input morphological parameters is analogous to a “performance landscape” (Simpson 1944; Niklas 1994; Arnold 2003), which illustrates the range of performance across a morphological parameter space. We calculate the ov erall kinematic performance as a weighted combination of the different kinematic output metrics in a manner similar to that used by (Stayton 2019). In that work, a combined metric was calculated as an arithmetic mean for each metric across a range of morphological parameters. For our context, in calculating a combined LaMSA ratio we use a geometric mean of the individual ratios,

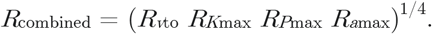

Using the geometric mean ensures that if the kinematic output of the LaMSA system was 100x better for one metric (*R* = 100) and 100x worse for another (*R* = 0.01), that the combined result would be *R* = 1 (an arithmetic mean would give *R* ≅50 in that case, misleadingly suggesting that the LaMSA system was overall 50x better).

## RESULTS & DISCUSSION

### Mechanical Sensitivity

Using the mean values of the focal morphological parameters from 12 *Strumigenys* taxa (Table 2), we calculated the sensitivity of the LaMSA and MDA models to these morphological parameters (see Methods: Model sensitivity analysis). Take-off velocity in the LaMSA model was most sensitive to changes in *F*_mus_ (Fig. 2A, left panel), while the take-off velocity of the MDA model was most sensitive to changes in MA (Fig. 2B, left panel). Sensitivity analyses of other kinematic output metrics such as KE_max_, *P*_max_, and *a*_max_ (Fig 2) shows that for the average *Strumigenys* measured, the model predicts that kinematic output is most sensitive to MA and *L*_mus_ for the MDA system while the LaMSA system is more sensitive to *F*_mus_, *M*_man_, and *L*_mus_.

**Figure 2:**
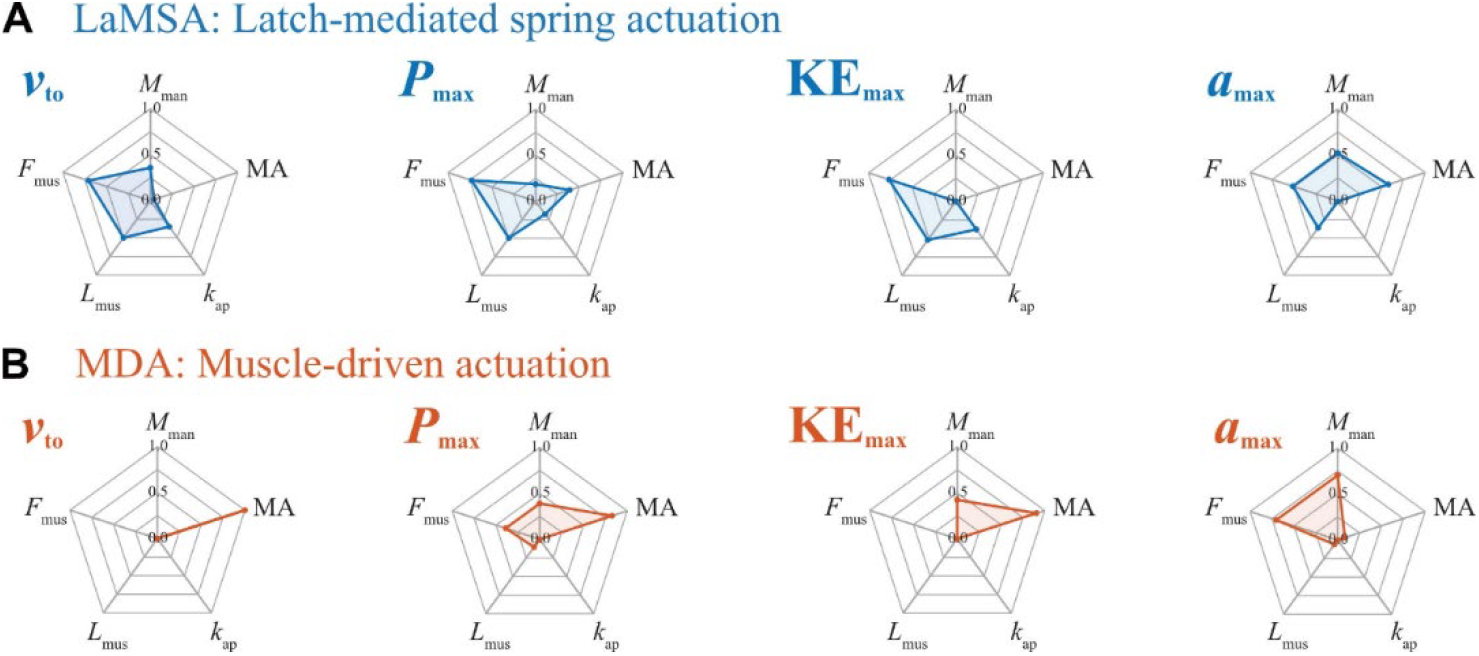
The kinematic outputs predicted to be most sensitive to different morphological parameters, compared across the LaMSA and MDA models. Spider web charts (radar charts) depict the sensitivity of output parameters to changes of each input parameter for both a LaMSA system (panel A) and an MDA system (panel B). For example, the take-off velocity (*v*_*to*_) of a LaMSA system (panel A, left column) is most sensitive to variation in muscle force and length (*F*_mus_ and *L*_mus_) and is least sensitive to variations in the MA of the mandible. For the MDA system, the take-off velocity (panel B, left column) is most sensitive to variations in mandible MA.

**Figure 3:**
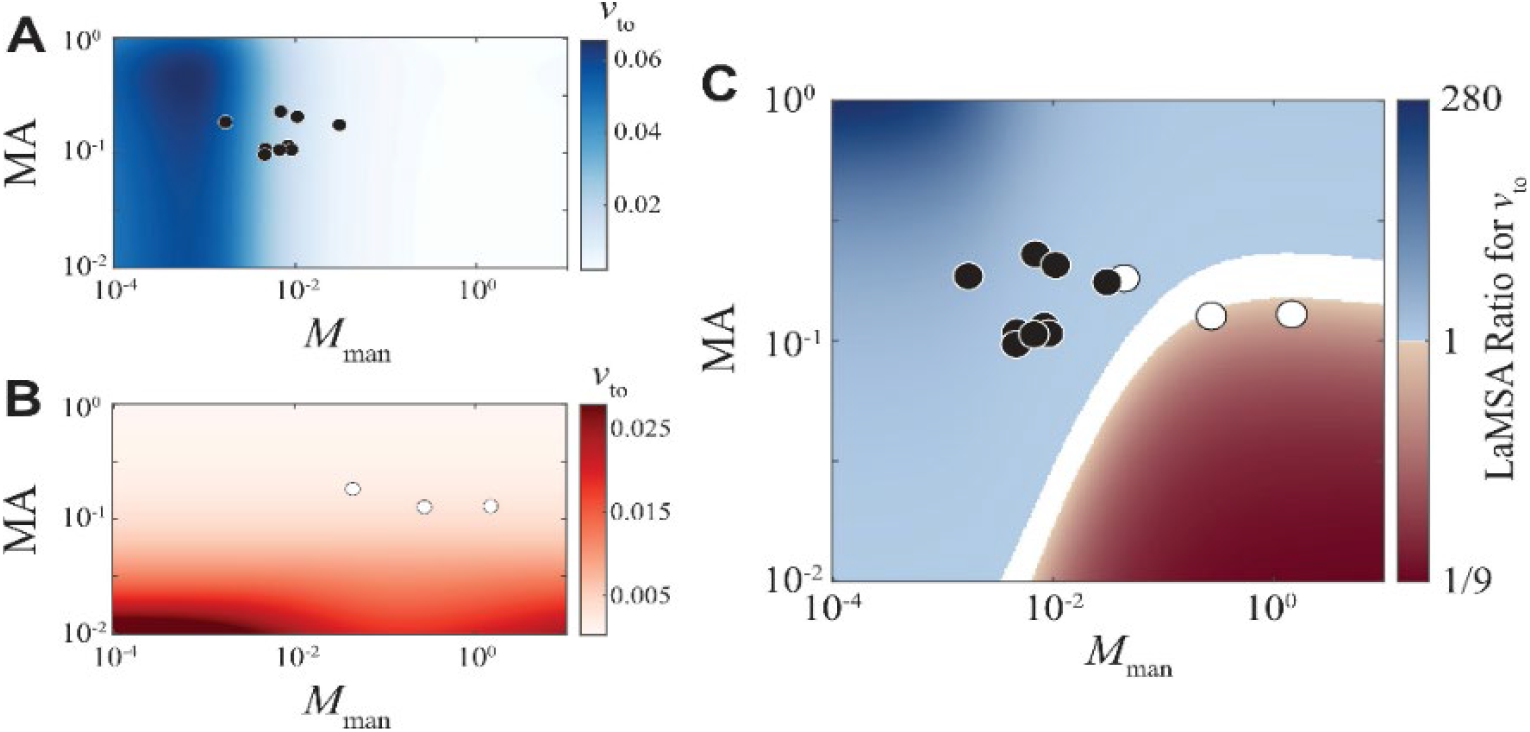
For input parameters representative of *Strumigenys*, the models predict that LaMSA outperforms MDA at small masses. **A**. The take-off velocity of the LaMSA system, where in this figure the apodeme spring constant, muscle length and its maximum force are “coupled” to the mandible mass: as ***M***_**man**_ increases on the x-axis, the values used for ***K***_**ap**_, ***L***_**mus**_, and ***F***_**mus**_ as inputs to the model also increase according to the equations from Table 3. The trap-jaw *Strumigenys* species are overlaid onto the morphospace (black circles). **B**. The take-off velocity of the MDA system for the same coupled morphospace as panel A with the non-trap jaw *Strumigenys* species overlaid (white circles). **C**. Taking the ratio of panel A/B, the LaMSA ratio for ***v***_**to**_ is largest at small values ***M***_**man**_. The LaMSA Zone (blue regions of the plot) corresponds to where a LaMSA system outperforms an MDA system with comparable morphology. White regions of the plot represent morphologies in which LaMSA and MDA result in similar performance (functional equivalence). The non-trap jaw (white circles) and trap-jaw (black circles) *Strumigenys* species are overlaid onto the morphospace.

**Figure 4:**
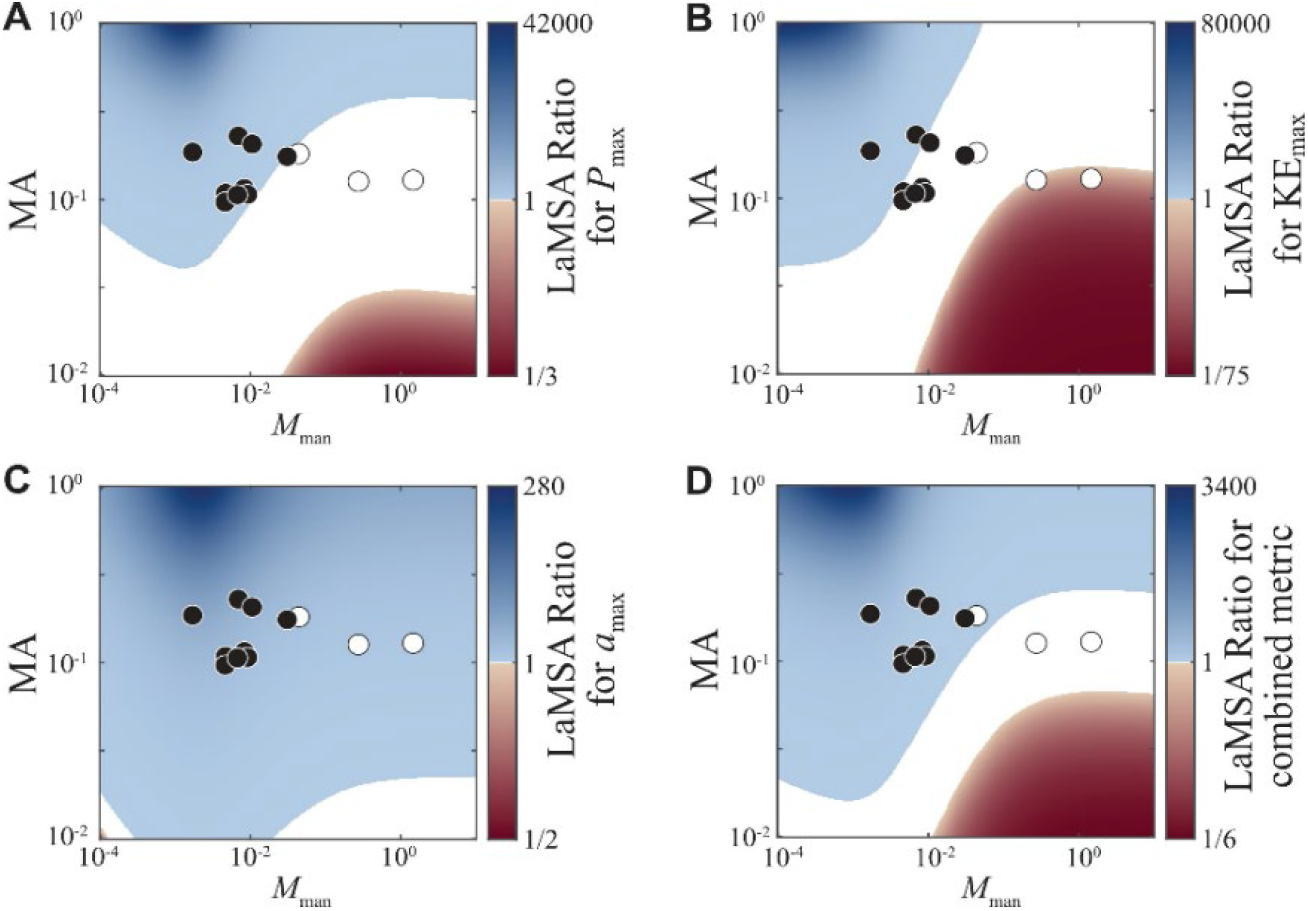
The size and shape of the LaMSA Zone can depend on the kinematic output metric. **A-C**. The LaMSA Ratio for a set of coupled input parameters is calculated for maximum power, maximum kinetic energy, and maximum acceleration. The LaMSA Zone (blue regions) and region of functional equivalence (white regions) occupy different amounts of the morphospace for each kinematic output metric. Compared to take-off velocity (Fig. 3C), ***P***_**max**_ displays a wider region of functional equivalence, and a much smaller LaMSA Zone in the explored parameter space. **D**. The ratio of the combined output at each point in the morphospace is calculated from a geometric mean of the take-off velocity, maximum power, maximum kinetic energy, and maximum acceleration.

Our first objective was to use our methodological framework to explore mechanical sensitivity in the trap-jaw LaMSA system. By analyzing performance gradients in 2D trait spaces we find that there is a great deal of variation in how changes in input variables influence different output variables across both LaMSA and MDA models (Fig. 2). Two major patterns emerge: *i*) Different kinematic output metrics show different patterns of mechanical sensitivity across the same set of input parameters. While velocity, power and kinetic energy are sensitive to muscle traits and insensitive to aspects of the mandible for the LaMSA system, acceleration is less sensitive to the muscle traits and does show sensitivity to mandible traits (Fig. 2). *ii*) Patterns of mechanical sensitivity are different between the LaMSA and MDA systems. While the LaMSA model generally shows a high sensitivity to muscle traits but less sensitivity to changes in the mandible; the MDA system shows high sensitivity to aspects of the mandible, and low sensitivity to muscle traits.

### Performance Transition

To simulate a performance space fully relevant to *Strumigenys*, we included the covariance between the different morphological parameters within the group (Table 3). The variance of MA among our measured species were uncorrelated with their measured *M*_man_, while the other three focal morphological parameters (*L*_mus_, *k*_ap_, *F*_mus_) were positively correlated with *M*_man_.

We used the observed correlations between morphological traits to define couplings between the model inputs. We created a coupled morphospace of take-off velocity for both LaMSA (Fig. 3A) and MDA (Fig. 3B) systems, where mandible mass was used to define the spring constant of the apodeme and muscle input through the correlations found in Table 3 (vertical slices in Fig. 3 of constant mandible mass also have fixed apodeme spring constant, muscle length, and maximum isometric muscle force). For the LaMSA system, there is a sharp decrease in the take-off velocity as a function mandible mass (Fig. 3A), which indicates that for the morphospace relevant to *Strumigenys* there is a maximum size for effective spring-driven movement. Interestingly, the trap-jaw *Strumigenys* species are clustered in the region of morphospace where the LaMSA system take-off velocity starts to decrease. For the MDA system, take-off velocity increases for smaller MA and larger mass (Fig. 3B).

To directly compare the two different categories of ants – trap-jaw vs. non-trap-jaw – we visualized regions where the LaMSA model outperformed the MDA model using the LaMSA ratio: the ratio of LaMSA kinematic output to MDA kinematic output. The LaMSA ratio metric allowed visualization of the relative kinematic performance of coupled parameter combinations (Fig. 3C). The LaMSA ratio for *v*_to_ partitioned the parameter space into two zones - one where LaMSA outperformed MDA and one where MDA outperformed LaMSA - separated by a boundary of equal output (Fig. 3C). We term this region the LaMSA zone, which approximately separated the two groups of *Strumigenys* based on their mandible actuation. Two of the three non-trap-jaw ant data points were within the region where the MDA model outperformed the LaMSA model, and all trap-jaw ant data points were within the region where LaMSA outperformed MDA. Notably, both trap-jaw and non-trap-jaw data points were close to the boundary of equal model output for *v*_to_.

When we examined the LaMSA Ratio for a different kinematic output metric, maximum power, we observed a LaMSA Zone at small mass and large MA, as we did for take-off velocity. In Figure 4A, the LaMSA Ratio for *P*_max_ is shown for the same coupled morphological inputs used in Figure 3C. The LaMSA Zone for *P*_max_ separated the two groups of *Strumigenys* based on their actuation type (Fig. 4A), with many of the species located close to the boundary of equal performance between LaMSA and MDA. This boundary of equal performance (the white regions of the plot) is significantly broader for *P*_max_ than it is for *v*_to_.

Calculating the LaMSA Ratio for other kinematic output metrics including maximum acceleration and maximum kinetic energy, revealed that the LaMSA zone for each metric had a different size in the parameter space (Fig. 4B-C). These metrics had a similarly shaped LaMSA zone as that of *v*_to_ and *P*_max_, but with slightly differing sizes. Maximum acceleration had the largest LaMSA zone, with the model predicting that a LaMSA system would outperform MDA for the full range of parameters explored.

To combine these four kinematic output metrics into a single combined morphospace, we applied a performance landscape analysis to our comparison of LaMSA versus MDA models for *Strumigenys* mandible closure using a combined metric (see Methods - Model comparison). We used the combined metric to visualize the effect of five parameters across four kinematic output metrics evaluated using a LaMSA model and an MDA model (Fig. 4D). Although each metric was equally weighted in the combined space, the size of the LaMSA zone for the combined space most closely resembled those for the plots of maximum power and take-off velocity.

The LaMSA zone plots illustrate a performance transition. These plots allow for direct comparison of multiple biomechanical models in a single morphospace, which shows regions of morphospace where the LaMSA and MDA models outperform each other and regions that result in equivalent performance (Fig. 3). The LaMSA zone plot is based on relative kinematic performance, delineating regions of input variance where the combination of morphological component values perform better as a LaMSA system (blue) and those regions where an MDA system would perform better (red). It also delineates a white region between the two, where the two biomechanical models show approximately equivalent kinematic performance. Interestingly, the size of this zone of kinematic equivalence varies a great deal, as seen when comparing the LaMSA zone plots for velocity and power output (Figure 3C and 4A). The equivalence zone for velocity is narrow, illustrating that there are few configurations of input parameters that result in similar velocity.

A final objective of our approach is to examine the performance transition across multiple kinematic output metrics by overlaying several performance gradients and using this to identify trade-offs between them. What we find is that there are regions of the stacked performance space where there wouldn’t necessarily be trade-offs. After overlaying gradients for four kinematic output metrics across our LaMSA space, there is a region where all four performance variables show local optima. Furthermore, this region is where the 9 measured trap-jaw taxa cluster when plotted in the morphospace. Another striking pattern is that for several kinematic output metrics, the three non-trap-jaw taxa fall within the white equivalence zones (Figures 3 & 4). This indicates that although they are using an MDA system, they could achieve similar values of power and kinetic energy to what they would achieve if they used a LaMSA system. Additionally, the different kinematic output metrics vary in their size of kinematic equivalence zones (white regions of Figs. 3-4). In contrast to the small equivalence zone for take-off velocity (Fig. 3C), the equivalence zone for power output is large (Fig. 4A), showing a great deal of equivalent configurations.

## Biological Relevance and Testable Predictions

### Mechanical Sensitivity

The variation in mechanical sensitivity across the kinematic output metrics could allow the morphology of the system to be altered in ways that change one metric without affecting another. A study performed on the complex cranial linkages connecting the jaws to the hyoid and suspensorium in fish skulls showed that it is possible to change particular bone lengths to alter jaw performance while allowing performance in the hyoid or suspensorium to remain unchanged (Baumgart and Anderson 2018). A similar situation could occur in the trap-jaw model seen here. Altering the mandibles of the LaMSA model should allow for shifts in acceleration without significantly altering other aspects of performance. Similarly, altering aspects of the spring could influence velocity and kinetic energy while leaving acceleration relatively untouched.

The mechanical sensitivity in these systems could also have implications for the evolution of trap-jaw and non-trap-jaw ants. Previous work on mechanical sensitivity in the linkage systems of both mantis shrimp appendages and fish jaws has suggested that the less sensitive performance is to change in a morphological component, the more free that particular component may be to diversify, as such changes will have less effect on potential fitness (P. S. L. Anderson and Patek 2015; Hu, Nelson-Maney, and Anderson 2017). Furthermore, those components that do have a strong influence on performance are prone to higher rates of evolution (Muñoz, Anderson, and Patek 2017; Muñoz et al. 2018). Given this, we might expect that the mandibles in *Strumigenys* trap-jaws should show wider diversification but lower evolutionary rates due to their lower influence on velocity, power, and energy output while the opposite would be true for the non-trap-jaw forms. Recent work examining the evolutionary rates of morphological characters in *Strumigenys* has shown that in some trap-jaw lineages, there is evidence for decreased evolutionary rates in the mandible relative to non-trap-jaw forms (P S L Anderson 2022). This same study also shows a slight increase in rates for traits related to the closer muscle size, which makes sense in light of the sensitivity of several kinematic outputs to muscle inputs in the LaMSA model (Figure 2).

The patterns of mechanical sensitivity for the trap-jaw system might be generalizable across other biological LaMSA systems. The higher sensitivity towards maximum isometric muscle force of the LaMSA versus MDA models of *Strumigenys* might be a general feature of biological LaMSA systems. For example, previous work found that mantis shrimp maximize muscle force through an increase in overall muscle physiological cross-sectional area (Blanco and Patek 2014) and frog species that use proportionally larger amounts of elastic energy in their jumps generate proportionally higher muscle forces (Mendoza and Azizi 2021). Based on our result s in *Strumigenys*, simulations of LaMSA versus MDA movements tuned to mantis shrimp or frogs may reveal a high sensitivity of LaMSA kinematic output to maximum isometric muscle force. That type of tuning could also be used to determine the overlap of different kinematic output metrics for other biological LaMSA systems.

While the mechanical sensitivity patterns identified here are intriguing and potentially give insights into the mechanics of the trap-jaw system, it is important to recognize a few caveats. First, we are exploring mechanical sensitivity for kinematic output metrics that are important to the trap-jaw ant but may not be as important to non-trap-jaws. Non-trap-jaw forms utilize their mandibles for static biting/gripping as opposed to kinetic strikes (Brown Jr and Wilson 1959). Therefore, kinematic variables may be less important than variables such as static bite forces, similar to what has been explored in other MDA insect mandibles (Goyens et al. 2014; Weihmann et al. 2015; David et al. 2016; Klunk et al. 2021; Püffel et al. 2021). Furthermore, we analyzed sensitivity as a given percentage change for each input, but some parameters might have larger variation than others, leading to different absolute changes. Both of these issues can be explored in future work with these models by examining a broader range of potential output variables and input variation.

### Performance Transition

Our work here extends ideas from previous work on LaMSA system modeling that found a mass-dependent transition between LaMSA and MDA performance (Galantis and Woledge 2003; Ilton et al. 2018). In that previous work, the question of whether a LaMSA system outperformed an MDA system was explored by fixing the motor and allowing the mass of the system to vary. Although that approach was useful to understand the kinematics of different actuation methods for a motor of a fixed size, it ignored the potentially large changes of motor properties when comparing across multiple systems within a taxonomic group. The LaMSA zone framework extends the single-variable mass-dependent transition between LaMSA and MDA performance into a multi-variable performance space. The many-dimensional parameter space that defines the LaMSA zone can be reduced to a lower-dimensional space using couplings between the input parameters, as we did here in the case of *Strumigenys* (we reduced a 5D space of focal parameters down to a 2D representation in Figures 3-4). Although this parameter reduction has similarities to previous LaMSA modeling that made isometric scaling arguments (Sutton et al. 2019; Hawkes et al. 2022), here our scaling relationships are specific to the taxa being investigated. In other words, we are tuning the models to the biological reality of these specific taxa.

The comparison between our tuned approach and previous work highlights a pit-fall of examining traits as independent entities in functional landscapes: such landscapes run the risk of deviating from biological reality. One of the strengths of the theoretical morphospace method for analyzing form-function relationships is being able to identify unoccupied regions of morphospace and examine why they aren’t occupied (Raup and Michelson 1965; McGhee 1999). The general assumption is that there will be some constraint, functional, structural, ecological, or developmental, preventing the theoretical morphologies within those regions from being realized in biology and identifying those constraints gives insights into the controls on morphological diversity. By ‘tuning’ the morphospace to our particular group of interest, we allow for a more precise analysis of the theoretical morphospace for *Strumigenys* and can query the unoccupied spaces knowing that they are possible but unrealized biological forms.

The modeling approach presented here allows for the incorporation of biological data which both makes the morphospace more biologically relevant and allows for the model to be scaled up with relative ease by adding more taxa. Tuning the morphospace to the taxa in question transforms it from a general morphospace to one tailored to the taxa being studied; a powerful tool for exploring the form-function relationship within a specific taxonomic group, allowing for the interdependence of morphological traits within a group to be accounted for. That said, it is important to remember that the relationship between traits is dependent on the taxa being measured. Our current case study only uses twelve taxa, so making broader claims about the group will require a larger dataset. Furthermore, our data comes from both trap-jaw and non-trap-jaw forms. It is unclear whether the different types would actually show significantly different scaling relationships between traits, which may complicate the analyses. However, the modeling approach is heavily adaptable, as adding or removing taxa is easily accounted for in a quantitative fashion. As taxa are added or removed, new correlations can be calculated and new tuned morphospaces can be created, leading to an even more biologically-informed performance landscape. Furthermore, our method can be extended to any group with a LaMSA system, or indeed any multi-part biomechanical system. All that is needed is a quantitative model incorporating the various input parameters and correlations of those parameters across the sample. Utilizing input correlations in this way can result in more focused studies of the form-function relationship within multi-part biomechanical systems.

It is rare that a biomechanical system only has a single measurable output metric. Even in the case of a simple lever system found in many vertebrate mandibles, it is possible to measure the force of a bite as well as its maximum speed, two metrics that generally form a trade-off in these systems (Westneat 2003). Trade-offs such as these are often viewed as potential drivers of speciation and thereby diversification due to creating multiple directions of selection (Taylor and Thomas 2014; Vincent 2016; Polly 2020; Waldrop et al. 2020). In multi-part systems such as LaMSA systems there are a variety of potential output metrics, all of which may be important functionally, such as velocity, acceleration, power, and energy. Previous work examining scaling in LaMSA models has shown that across a wide diversity of forms, the specific optima for these metrics are often not concurrent (Ilton et al. 2018). There is some overlap between LaMSA zones for different metrics, but this overlap is incomplete. Our results seem to show that the trap-jaw ants are centered on a region of morphospace where the LaMSA zones for all four metrics overlap. While intriguing, a larger diversity of taxa will be required to test whether this pattern holds true across the wide diversity of *Strumigenys* LaMSA forms (Booher et al. 2021). The methods presented here can accommodate such an increase in data and could be used to test for patterns of LaMSA zone occupation and potential morpho-functional constraints arising from conflicting performance optima.

The biological significance of the three non-trap-jaw taxa in the equivalence zone is harder to interpret. While it seems intuitive that they should fall in the MDA zone, it is necessary to remember that the specific performance measures being compared are kinematic measures, which may not be what drives non-trap-jaw evolution. More likely, static bite force or fine-scale manipulation would be the driving functions for non-trap-jaws, but those performance metrics are not included here. That said, it should be noted that a larger kinematic equivalence zone may allow these taxa the freedom to shift their morphology to better optimize the system for more static performance, while maintaining a certain level of kinematic performance. Future work can easily add or subtract performance metrics depending on the question and taxa being analyzed, allowing for direct testing of different performance metrics and ecologies.

## CONCLUSION

Here we examined core principles of form-function relationships in complex, multi-part biological systems. While we utilized the LaMSA system in a genus of trap-jaw ants as our study mechanism, this framework can be used for any sufficiently complex system. All that is required is a working model of the system as well as some data on the variation in morphological components. These can then be used to construct a morphospace tuned to the system via trait correlations along with kinematic performance gradients derived from the model. If several alternative models for the function of a system exist, these can be incorporated and directly compared via the LaMSA zone method. Altogether, this framework allows for the non-linearity in form-function relations to not just be accounted for, but directly examined and tested within these systems.

Biomechanical models are often used to explore non-linear relationships between form and performance. However, these models must walk a fine line between biological reality and computational pragmatism. The urge is always to make the models as close to reality as possible; however, this can lead to models that are too complex to be solved and of limited use outside of individual systems. There is value in utilizing more generalized ‘simple’ models which have broader application and reduced computational load (Philip S L Anderson, Rivera, and Suarez 2020). However, generalized models can run afoul of examining aspects of morphological or performance space that are simply irrelevant to biology. This balance between simplicity and biological relevance is similar in nature to the templates and anchors paradigm (Full and Koditschek 1999), where a simplified template model describes general biomechanical principles and an anchor model explores specific realizations of the more general principle. Intermediate to these two extremes, in this work we presented a method for incorporating biological data into a generalized mathematical model of a LaMSA system. This method allows for tuning a generalized model, which is easier to manipulate and explore, to specific biological systems.

## ACKNOWLEDGEMENTS

The authors thank S. N. Patek for contributions to the ideas in this work. M.I. acknowledges funding support from the NSF for this work under grant no. 2019371 and P.S.L.A. acknowledges funding from NSF IOS 17-55336. J. T. C., R. L. D. and A. C. acknowledge funding support from the Harvey Mudd College Physics Summer Research Fund.

## Notes

### Competing Interest Statement

The authors have declared no competing interest.

